# Targeting ZNF638 activates antiviral immune responses and potentiates immune checkpoint inhibition in glioblastoma

**DOI:** 10.1101/2024.10.13.618076

**Authors:** Deepa Seetharam, Jay Chandar, Christian K. Ramsoomair, Jelisah F. Desgraves, Alexandra Alvarado Medina, Anna Jane Hudson, Ava Amidei, Jesus R. Castro, Vaidya Govindarajan, Sarah Wang, Yong Zhang, Adam M. Sonabend, Mynor J. Mendez Valdez, Dragan Maric, Vasundara Govindarajan, Sarah R. Rivas, Victor M. Lu, Ritika Tiwari, Nima Sharifi, Emmanuel Thomas, Marcus Alexander, Catherine DeMarino, Kory Johnson, Macarena I De La Fuente, Ruham Alshiekh Nasany, Teresa Maria Rosaria Noviello, Michael E. Ivan, Ricardo J. Komotar, Antonio Iavarone, Avindra Nath, John Heiss, Michele Ceccarelli, Katherine B. Chiappinelli, Maria E. Figueroa, Defne Bayik, Ashish H. Shah

## Abstract

Viral mimicry refers to the activation of innate anti-viral immune responses due to the induction of endogenous retroelement (RE) expression. Viral mimicry has been previously described to augment anti-tumor immune responses and sensitize solid tumors to immunotherapy including colorectal cancer, melanoma, and clear renal cell carcinoma. Here, we found that targeting a novel, master epigenetic regulator, Zinc Finger Protein 638 (ZNF638), induces viral mimicry in glioblastoma (GBM) preclinical models and potentiates immune checkpoint inhibition (ICI). ZNF638 recruits the HUSH complex, which precipitates repressive H3K9me3 marks on endogenous REs. In GBM, ZNF638 is associated with marked locoregional immunosuppressive transcriptional signatures, reduced endogenous RE expression and poor immune cell infiltration (CD8^+^ T-cells, dendritic cells). ZNF638 knockdown decreased H3K9-trimethylation, increased cytosolic dsRNA and activated intracellular dsRNA-signaling cascades (RIG-I, MDA5 and IRF3). Furthermore, ZNF638 knockdown upregulated antiviral immune programs and significantly increased PD-L1 immune checkpoint expression in patient-derived GBM neurospheres and diverse murine models. Importantly, targeting ZNF638 sensitized mice to ICI in syngeneic murine orthotopic models through innate interferon signaling. This response was recapitulated in recurrent GBM (rGBM) samples with radiographic responses to checkpoint inhibition with widely increased expression of dsRNA, PD-L1 and perivascular CD8 cell infiltration, suggesting dsRNA-signaling may mediate response to immunotherapy. Finally, we showed that low ZNF638 expression was a biomarker of clinical response to ICI and improved survival in rGBM patients and melanoma patients. Our findings suggest that ZNF638 could serve as a target to potentiate immunotherapy in gliomas.

## INTRODUCTION

Glioblastoma (GBM) is the most common primary brain tumor in adults with poor median survival rates that have minimally changed over the last 20 years.^1^ Due to failures in the current standard-of-care treatment regimens, Immune Checkpoint Inhibition (ICI) has been proposed for high-grade gliomas given their success in other solid tumors.^2^ However, clinical trials using ICI for gliomas have largely failed due to i) poor tumor antigen presentation ii) scant intratumoral lymphocyte infiltration iii) reduced immune checkpoint presentation, iv) poor solid tumor penetration of the immunotherapy and v) immune-silencing via myeloid-derived suppressor cells/microglia.^3,4^ Therefore, strategies to overcome the constitutive immunosuppressive tumor microenvironment of GBM remain vital to improving outcomes for immunotherapy for GBM. Recently, viral mimicry has been proposed as a strategy to overcome the immunosuppressive tumor microenvironment by activating anti-viral anti-tumor immune responses.^5^ Viral mimicry refers to an activated antiviral cellular state that is triggered by the epigenetic activation of endogenous nucleic acids, often from retrotransposable elements (cytosolic dsRNA and DNA).^6,7^ Forming more than 40% of our human genome, retrotransposons such as Human Endogenous Retroviruses (HERVs) or LINEs (Long Interspersed Nuclear Elements) are normally silenced by epigentic modifications such as DNA hypermethylation, chromatin remodeling, and histone modifications.^4,5^ However, epigenetic dysregulation, such as the global DNA/histone demethylation found in GBM, facilitates reactivation of these viral-like sequences, inducing endogenous interferon responses mediated by innate dsRNA sensing pathway (RIG-I and MDA5).^6,7^ Viral mimicry has been used to potentiate tumor cytotoxicity and induce pre-clinical responses to immunotherapy in a variety of cancers, including colorectal cancer, melanoma, lymphoma, ovarian, and renal cell carcinoma.^8–11^ Since ICI trials in GBM have largely failed, investigating the role of epigenetic reprogramming and associated viral mimicry-induced immune responses presents an opportunity to enhance the efficacy of ICI.

Histone modifications, specifically H3K9me3, are the predominant epigenetic regulatory mark of retrotransposons. One of the main regulatory mechanisms of H3K9-mediated repression of endogenous retroelements occurs via the HUSH complex.^12^ The Human Silencing Hub (HUSH) complex is well conserved in mammalian genomes as a host defense mechanism against retroelements. Throughout evolution, the HUSH complex has not only protected against exogenous retroviruses (Human Immunodeficiency Virus, Murine Leukemia Virus), but also against retrotransposition of endogenous retroelements including HERVs and LINE-1 elements.^13^ The HUSH complex is composed of MPP8 (M-Phase Phosphoprotein 8), TASOR (Transcription Activation Suppressor), PPHLN1 (Periphilin 1) which recruits a histone methyltransferase, SETDB1 SET Domain Bifurcated Histone Lysine Methyltransferase 1 (SETDB1).^14^ A DNA binding protein ZNF638 has recently been found to be essential in the recruitment of HUSH, which forms the retroviral silencing complex (RSC) to ultimately precipitate H3K9me3 repressive epigenetic marks on unintegrated retroviral DNA.^12^ When this mechanism is lost in tissues, H3K9me3 repressive marks are removed, and cells become susceptible to endogenous retrotransposition, increasing HERV and LINE-1 expression and ultimately antiviral immune responses.^12,13^ **Figure 1A**. This viral mimicry effect has been demonstrated in other tumors where retroelements increased anti-tumor immunity and sensitized tumors to ICI.^15,16^ Here, we investigated the role of epigenetic reprogramming and associated viral mimicry-induced immune responses to enhance the efficacy of ICI in GBM through a novel epigenetic regulator, ZNF638.

**Figure 1.**
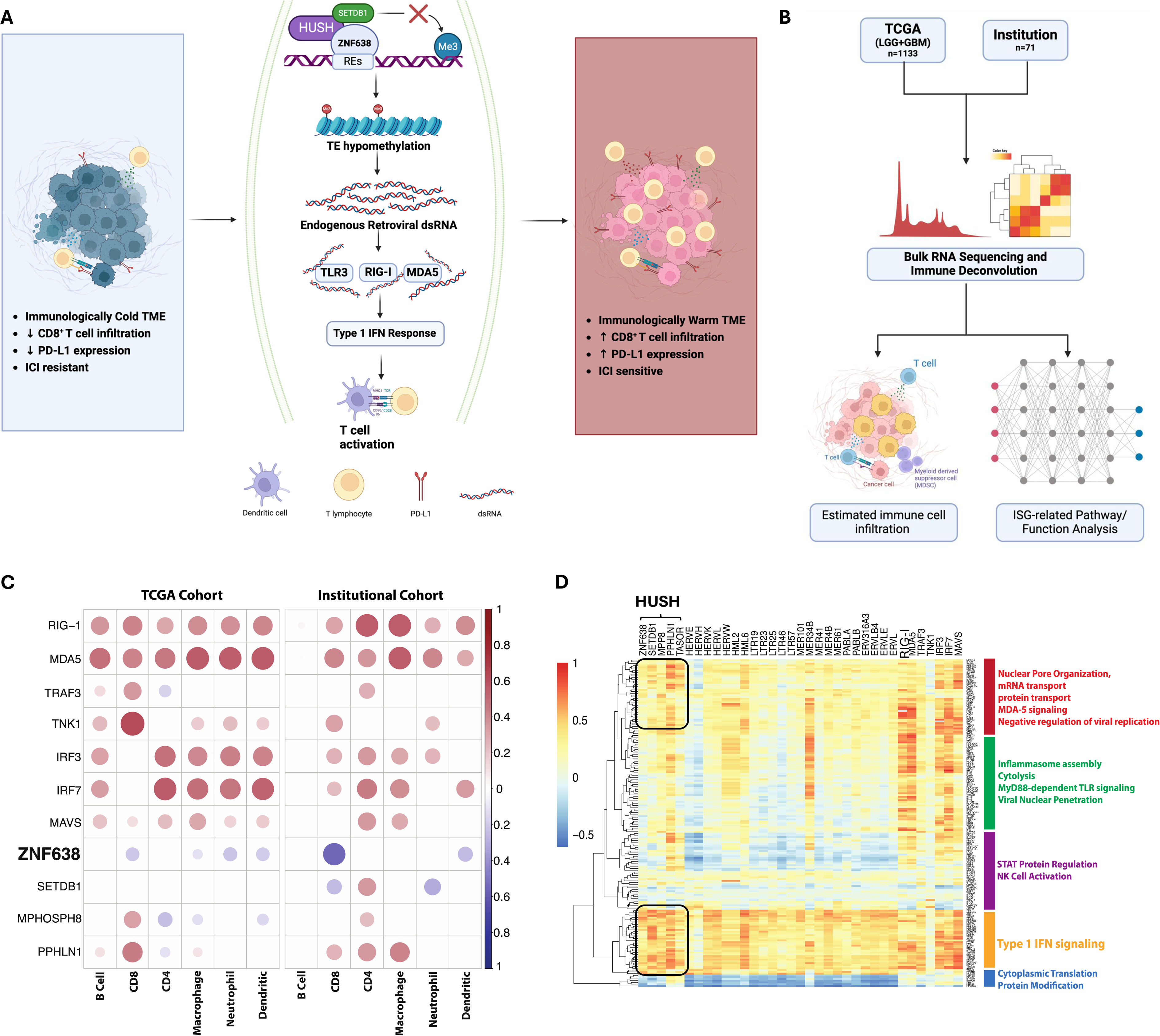
The retroviral silencing complex mediates the suppression of immunogenic RNA species in gliomas. A) ZNF638 acts as the master regulator of a retroviral silencing complex to silence retroelement expression via H3K9 trimethylation. Removal of HUSH-mediated repressive marks enhances anti-viral immune responses through innate dsRNA signaling. Made with BioRender. B) Bulk RNA sequencing data from TCGA GBM (n=617) and LGG (n=516) cohorts and our institutional cohort (n=71) were analyzed to conduct immune deconvolution and assess interferon-stimulated gene (ISG) related pathways and functions. Made with BioRender. C) The HUSH complex and ZNF638 are inversely correlated with CD8 immune cell infiltration (R_TCGA_= –0.2017, R_inst_=-0.5409) based on data obtained from TCGA GBM (n=617) and LGG (n=516) database and an institutional cohort (n=71). D) Correlation matrix demonstrates enrichment of ISGs with expression of several REs as well as a negative association between the HUSH complex and MDA5 signaling using the Reactome Pathways database. Gene ontology analysis demonstrates ZNF638 and HUSH complex are directly correlated with increased inhibition of NK cell activation and Type 1 IFN signaling. IFN=Interferon. TE = transposable elements.

## RESULTS

### ZNF638 is associated with a unique epigenetic and immunological landscape in GBM

Given the established role of ZNF638 in epigenetic silencing of retroelements, we sought to understand its association with dsRNA-sensing pathways in GBM. **(Figure 1A, Supplementary Figure 1 A,B).** To gain insight into the role of ZNF638 in shaping the epigenetic and immunological characteristics of GBM, we performed immune deconvolution using RNA sequencing from both The Cancer Genome Atlas (TCGA) and validated results in an independent institutional cohort of high-grade gliomas **(Figure 1B).** ZNF638 was significantly negatively correlated with dsRNA sensing pathways (RIG-I, MDA5, TLR3) in GBM and positively associated with members of the HUSH complex and its effectors (SETDB1 and MPHOSPH8) **(Supplementary Figure 1C).** Leveraging the Tumor Immune Estimation Resource (TIMER)^17^, we evaluated the association between the HUSH complex and the RIG-I sensing pathways with tumor lymphocyte infiltration. ZNF638 was strongly negatively correlated with CD8 T-cell infiltration in two independent datasets. Conversely, the viral mimicry cascade is also strongly correlated with immune cell infiltration (CD8, CD4, DC, NK, Macrophages). (**Figure 1C)** To recapitulate these findings, we demonstrated that ZNF638 exhibited a positive correlation with the HUSH components and a negative correlation with dsRNA sensing programs in brain tumor tissue based on data derived from the ARCHS4 dataset. These correlations are maintained when visualizing genome wide co-expression. ^18,19^**(Supplementary Figure 2A).**

To understand the landscape of endogenous retroviruses in GBM and their association with the HUSH complex, we utilized a custom bioinformatic pipeline using Telescope.^20^ This comprehensive analysis included the expression of 48 HERV families and the components of the HUSH/ZNF638 complex. Our findings indicated that the expression of HUSH effector proteins, PPHLN1, and TASOR exhibited widespread negative associations with several HERV families **(Supplementary Figure 2B).** Furthermore, using hierarchical clustering from human GBM specimens, we discovered that ZNF638 expression was associated with the downregulation of MDA5 signaling, while multiple HERV loci were enriched in the Type 1 Interferon cellular signaling programs **(Figure 1D).** Utilizing spatial transcriptomics, we demonstrated that regions with high expression of ZNF638 demonstrate upregulated transcription of components of the IFN-α and NFκβ signaling pathways as annotated by the Molecular Signatures Database (MSigDb) **(Figure 2)**. ^21^

**Figure 2.**
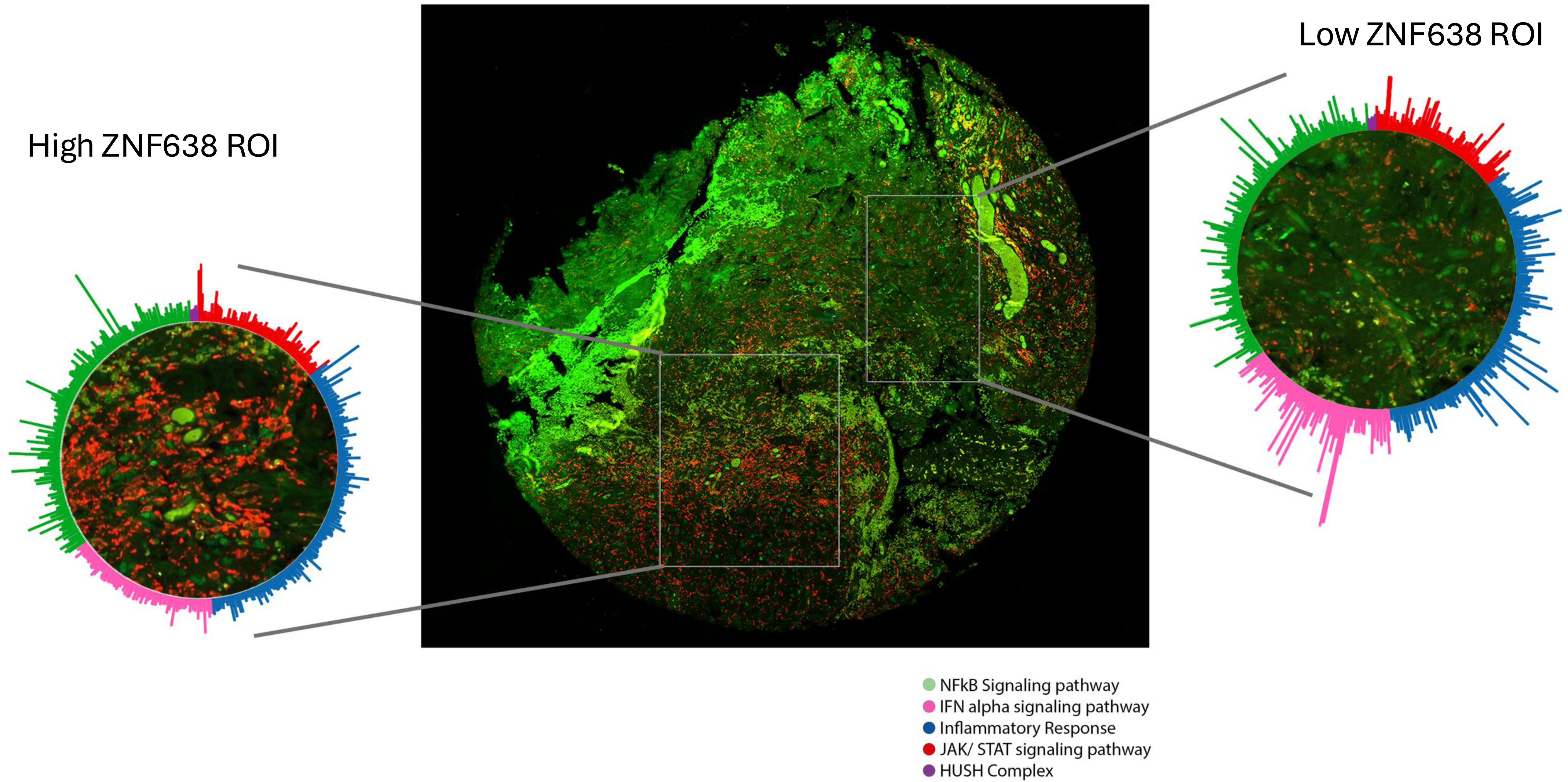
dsRNA-sensing and interferon signaling pathways are transcriptionally upregulated in high ZNF638 regions Circular visualization of gene expression in regions of high and low ZNF638 from a patient sample diagnosed with IDH WT recurrent GBM. Gene expression involved for the NFkB signaling pathway (n_genes_=200), IFN alpha signaling pathway (n_genes_=97), inflammatory response (n_genes_=200), JAK/STAT signaling pathway (n_genes_=87), and the HUSH complex (n_genes_=200), are shown. All gene sets obtained from the molecular signatures database (MSigBr). Central image shows the full tumor sample from the patient. Red: CD45, Green: Olig2

To understand the role of ZNF638 and the HUSH complex in GBM cellular states and the tumor environment, we leveraged a single-cell RNA sequencing dataset of 11 adult glioma patients to evaluate the impact of ZNF638 expression on expression of HERV families and associated dsRNA sensing pathway components.^22^ Retroelements were characterized in this dataset using TE-transcripts to create a custom dataset with a robust representation of the transcriptome and retrotranscriptome in glioma. Malignant cells lacking ZNF638 expression were significantly enriched in total retroelement expression (both coding and non-coding RNA elements). ZNF638 expression was associated with distinct cell-state clustering in oligodendrocyte-progenitor-like and neural-progenitor-like Neftel states. **(Figure 3A)** Cellular transcription states have been characterized as oligodendrocyte-progenitor-like (OPC, high PDGFR), astrocyte-like (AC, high EGFR), mesenchymal-like (MES, NF1 alteration), and neural-progenitor-like (NPC, high CDK4).^23^ Malignant cells with low ZNF638 expression demonstrated increased expression of total retroelements and increased expression of dsRNA sensing pathway regulators (RIG-I, TLR3, MAVS, MDA5, IRF3, IRF7). **(Figure 3B)** Furthermore, when stratifying individual tumors based on ZNF638 expression, we identified that low ZNF638-expressing tumors were associated with increased tumor-infiltrating lymphocytes. **(Figure 3C)** Differential infiltration of specific lymphocytes and cell types demonstrated a trend of increased infiltration of B-cells, Myeloid Cells, T cells, and Oligodendrocytes; however, p-values were >0.05 due to the small total number of immune cells. **(Figure 3D)**

**Figure 3.**
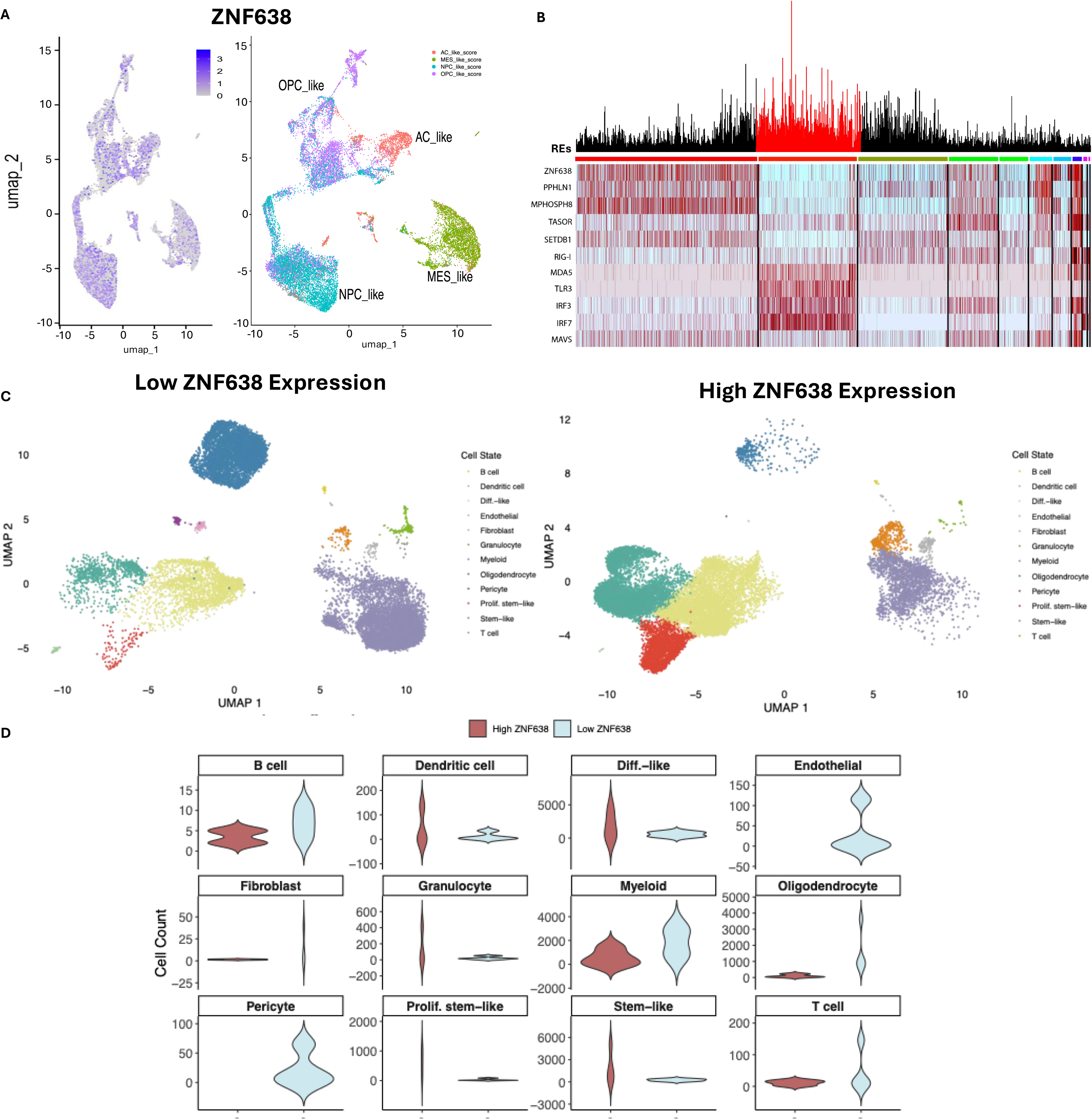
ZNF638 suppresses total RE expression and dsRNA sensing. A) Single-cell RNA sequencing and clustered analysis demonstrate that low ZNF638 is associated with enriched expression of total retroelements (REs) (n=18,400 cells, REs=5,680). B) Reduced cellular expression of the retroviral silencing complex (ZNF638, SETDB1, PPHLN1, MPHOSPH8, TASOR) is associated with increased total retroelement expression and increased expression of genes involved in the dsRNA sensing pathway (DDX58, IFIH1, TLR3, IRF3, IRF7, MAVS) (n=18,400 cells, REs=5,680). C) Low ZNF638 expression is associated with increased lymphocyte expression in individual GBM tumors using unsupervised clustering. UMAPs from tumors with low (n_tumors_=4, n_cells_=17535) and high (n_tumors_=4, n_cells_=20517) ZNF638 expression show heterogenous and distinct enrichment of cell types. D) Violin plots illustrate the expression levels of B cells, dendritic cells, differentiated-like cells, endothelial cells, fibroblasts, granulocytes, myeloid cells, oligodendrocytes, pericytes, proliferative stem-like cells, stem-like cells, and T cells in tumors with low versus high ZNF638 expression, based on unbiased cell type annotation. While a trend toward increased infiltration of T cells, B cells, myeloid cells, and oligodendrocytes is observed, this did not reach statistical significance likely due to the limited number of samples. Single cell data for panels A-D obtained from the European Genome-phenome Archive under accession number EGAS00001005300.

To confirm the clinical relevance of ZNF638 in GBM, we demonstrated that ZNF638 is uniquely enriched in GBM tissue relative to matched normal cerebral cortex as demonstrated by immunohistochemistry and western blot. **(Figure 4A, B, C)**. Using data from the National Cancer Institute Clinical Proteomic Tumor Analysis Consortium (CPTAC) we further demonstrate ZNF638, TASOR, MPHOSPH8, SETDB1 proteins are significantly enriched in GBM vs. normal brain tissue. **(Figure 4D, Supplementary Table 1)**

**Figure 4.**
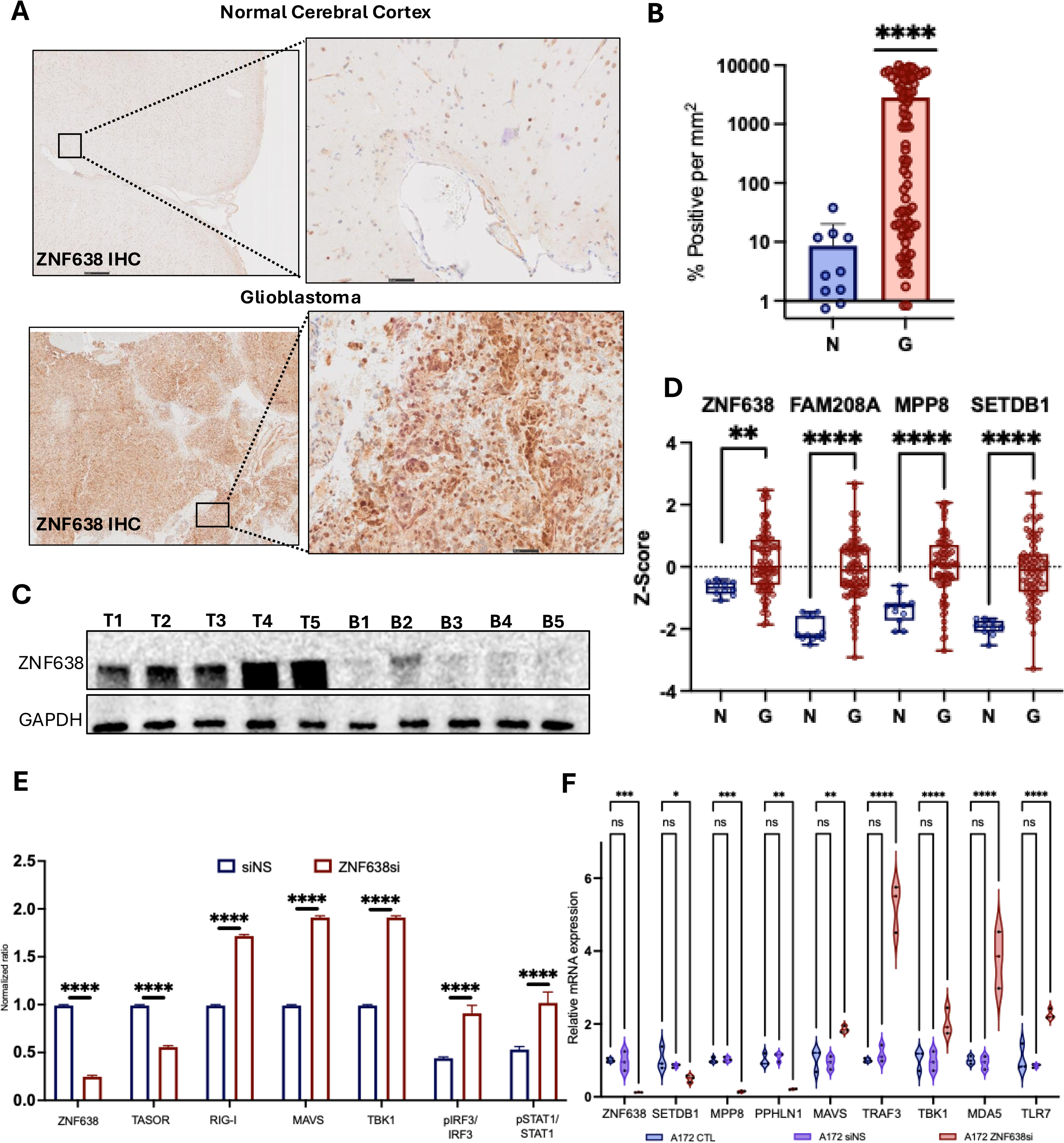
ZNF638 expression is enriched in GBM and induces dsRNA signaling when knocked down. A,B) Immunohistochemistry staining and quantification for ZNF638 demonstrates marked overexpression of ZNF638 in GBM tumors compared to matched normal cortex (n=48 vs. 10, p<0.0002). Results independently verified in biological replicate from two separate patient cohorts. C) Western Blot demonstrates that ZNF638 expression is enriched in GBM tumor samples compared to patient matched adjacent normal brain (n_T_=5 vs. n_C_=5). D) Proteomic data from the Clinical Proteomic Tumor Analysis Consortium (CPTAC) data portal for GBM (n=12) and normal tissue (n=99) corroborates significant enrichment of ZNF638, TASOR (FAM208A), MPHOSPH8, and SETDB1 in tumor tissue. E) Western Blot quantification validates that ZNF638 transient KD by siRNA reduces expression of HUSH via MPP8 and increases expression of RIG-I, MAVS, TBK1, pIRF3, and pSTAT1 in patient-derived GBM43 (ANOVA, ****P < 0.0001, ***P < 0.001, **P < 0.01). F) Knockdown of ZNF638 by siRNA decreases expression of SETDB1, PPHLN1, and MPP8 as well as increases expression of MAVS, TRAF3, TBK, MDA5, and TLR7 as measured by qPCR (biological triplicates, ANOVA, ****P < 0.0001, ***P < 0.001, **P < 0.01).

### ZNF638 Knockdown Suppresses HUSH Expression and Induces Innate Antiviral Immune Signaling

Given these findings, we sought to decipher the role of ZNF638 in regulating anti-viral immune signaling via epigenetic reprogramming of the retroviral silencing complex (RSC). To assess the role of ZNF638 in mediating the HUSH complex and dsRNA signaling pathway, we transiently downregulated ZNF638 in diverse GBM cell lines using RNA interference. Knockdown of ZNF638 resulted in a notable reduction in the expression of the RSC complex including SETDB1, PPHLN1, and MPP8 and increase in the innate antiviral signaling cascade, including MAVS, TRAF3, TBK1, pIRF3, and TLR7. **(Fig. 4E, F).**

To understand the molecular underpinnings of the viral mimicry cascade, we sought to understand the relationship between ZNF638, H3K9 trimethylation and dsRNA expression. Using dsRNA immunoprecipitation (J2 antibody, we demonstrated that ZNF638 knockdown significantly increased dsRNA expression in GBM A172 cells. **(Figure 5A-C).** Specifically, we found broadly increased expression of retroelements including LINE-1, Alus and LTRs with relatively no change in HERV-K RNA species. **(Figure 5B)**. This global increase in dsRNA intermediates was secondary to a reduction in H3K9 trimethylation. To confirm this result, we validated these findings using quantitative immunofluorescence and flow cytometry. Importantly, this viral mimicry activation was associated with an increase in immune checkpoint presentation (PD-L1) in GBM cells, as demonstrated by immunofluorescence, western blot and flow cytometry. **(Fig. 5A, C).** Additionally, ZNF638 knockdown induced downstream antiviral dsRNA signaling as evidenced by phospho-interferon regulatory factor 3 (pIRF3) expression on immunofluorescence. **(Figure 5D)** Using co-immunoprecipitation, we demonstrated that knockdown of ZNF638 resulted in significantly decreased levels of TASOR, MPP8, and SETDB1 in GBM cells, suggesting that ZNF638 is critical in maintaining the integrity of the HUSH complex and mediating H3K9me3 repressive histone marks **(Figure 5E, Supplementary Figure 3B).** In combination, these results signify that ZNF638-KD neutralizes HUSH-mediated H3K9 trimethylation in GBM and results in increased dsRNA signaling to stimulate the innate antiviral immune response.

**Figure 5.**
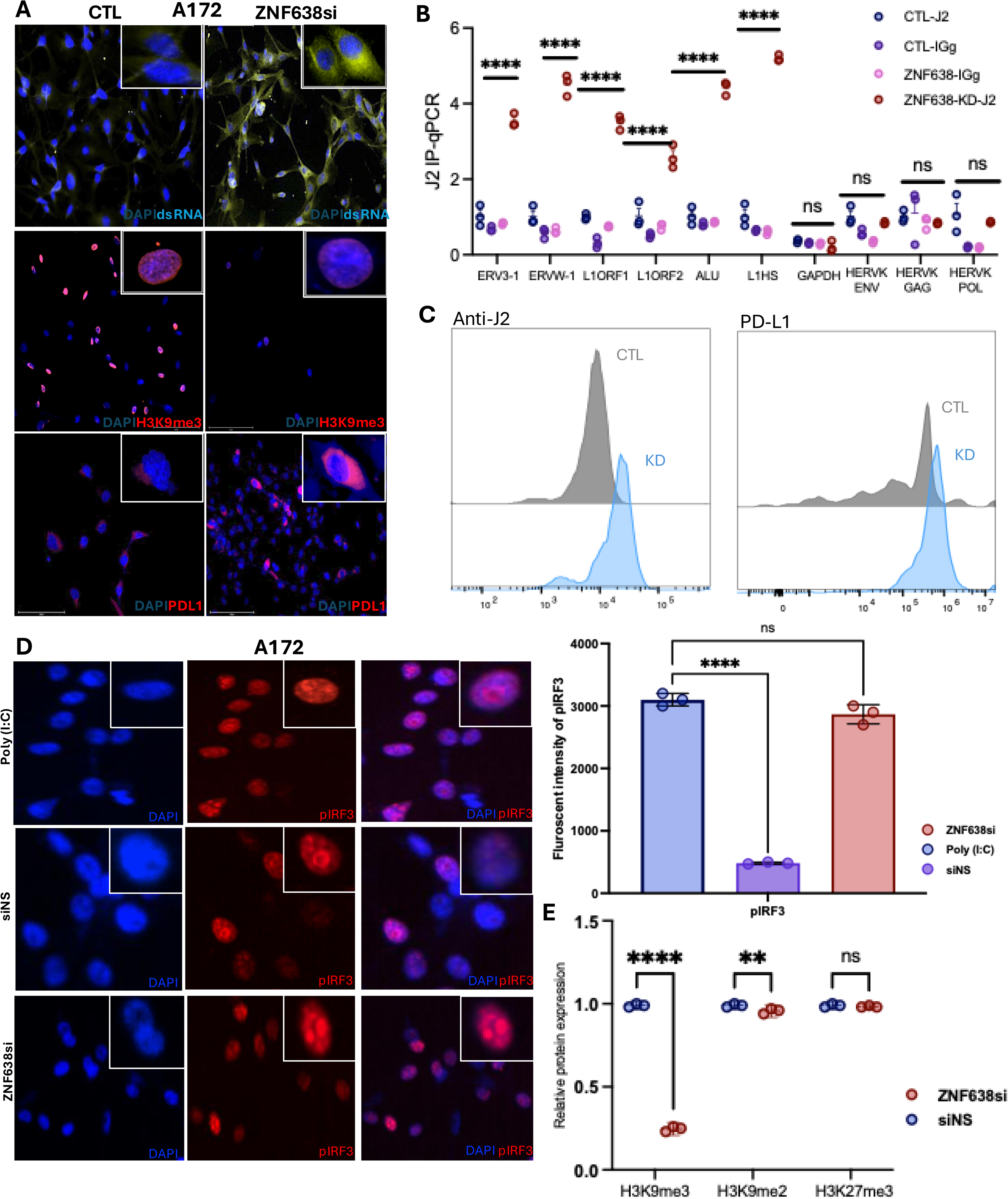
ZNF638 KD increases dsRNA expression secondary to loss of H3K9me3 signature. A) ZNF638 knockdown by siRNA in A172 GBM cells demonstrates increased expression of dsRNA (yellow), decreased H3K9me3 (red), and increased expression of PD-L1 (red) as evidenced by quantitative immunofluorescence. Nuclei are stained blue with DAPI. B) RNA-immunoprecipitation (J2 antibody) demonstrates increased pulldown of RE dsRNA with ZNF638 KD in A172 (performed in biological triplicate, ANOVA, **** P < 0.0001). C) Flow cytometry demonstrates increased expression of dsRNA (anti-J2) and PD-L1 with ZNF638 KD. D) ZNF638 knockdown elicits antiviral immune signaling via increased expression of pIRF3 (red) via quantitative immunofluorescence. Poly I:C (40ng/ml) represents a positive control for pIRF3 signaling. Nuclei are stained blue with DAPI. (ANOVA, **** P < 0.0001). E) Knockdown of ZNF638 with siRNA results in loss of H3K9me3 in A172 based on Western Blot (performed in biological triplicate, **** P < 0.0001, *P<0.05). ZNF638-KD does not change H3K27 trimethylation.

To decipher the role of ZNF638 in epigenetic and transcriptomic signatures in primary GBM neurospheres, we utilized a multi-omic bioinformatic approach using ATAC-seq and RNA-seq in patient-derived neurospheres. **(Figure 6A, B).** Using ATAC-seq, we noted that ZNF638 KD resulted in global epigenetic changes with increased open chromatin around select retrotransposons such as Tigger15a and AluJb **(Figure 6A)** Similarly, ZNF638 inhibition significantly upregulated innate immune and anti-viral programs. **(Figure 6C).** Specifically, ZNF638-KD significantly elevated intronic and retroelement transcripts globally with upregulation of specific retrotransposons (LTRs, LINEs, and Tigger1) utilizing a custom bioinformatics pipeline for retrotransposons. **(Supplementary Figure 4, Figure 6D, Supplementary Table 2, 3)**

**Figure 6.**
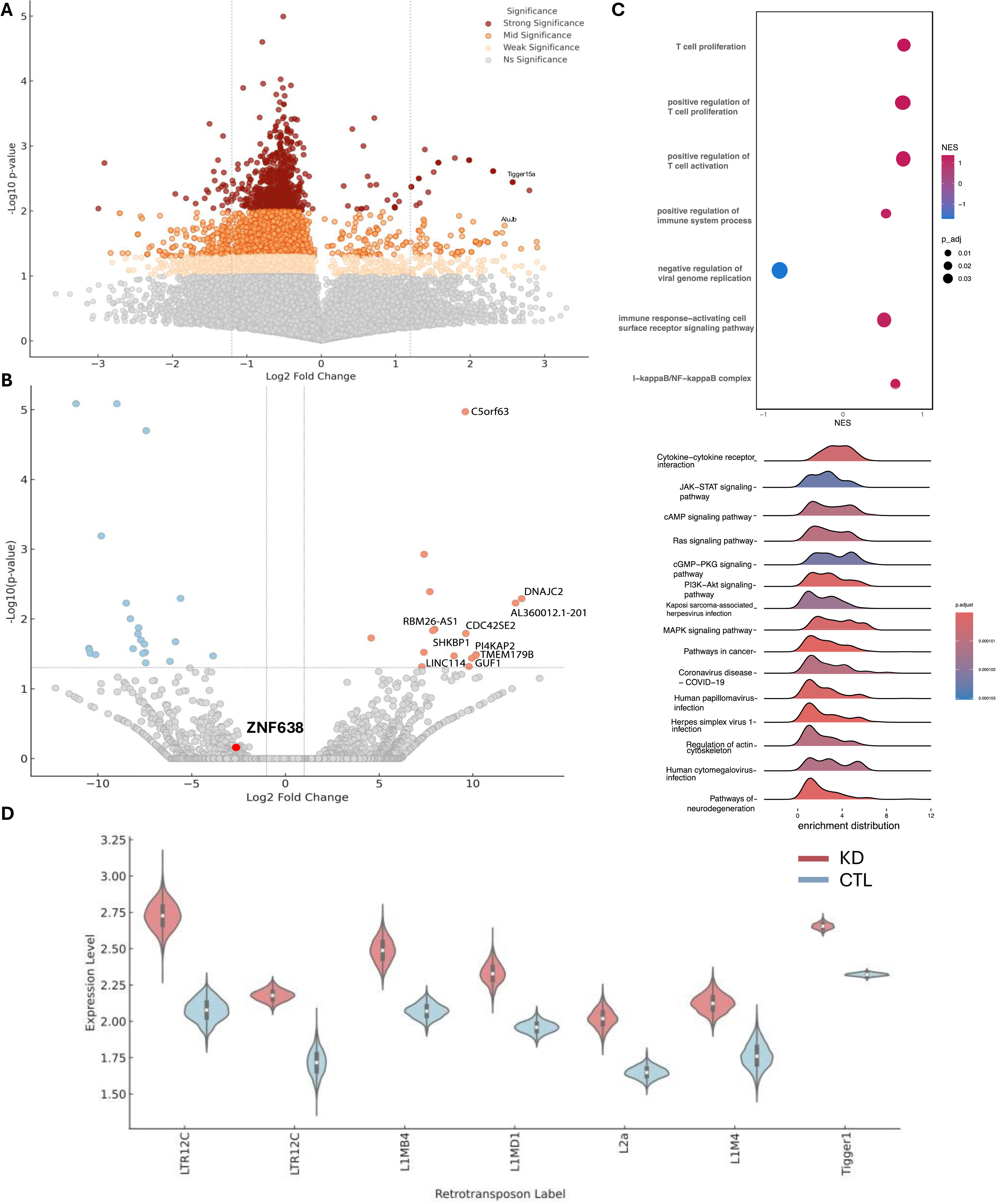
ZNF638 knockdown upregulates retroelement expression and corresponding antiviral and immune programs. A) Volcano plot shows differential analysis of ATAC-sequencing and demonstrates that ZNF638 KD in a patient-derived GBM cell line results in opening of genomic regions associated with repeat elements (AluJb, Tigger15a) (log_2_fold change_AluJb_=2.69 p_AluJb_=0.038, log_2_fold change_(Tigger16A)_=2.57 p_Tigger16A_=0.036). B) Volcano plot of differentially expressed genes from a patient-derived GBM cell line shows upregulation of transcripts related to the innate immune system (TMEM179B,log_2_fold change=10.1, p = 0.033), actin regulation in activated T-cells (CDC42SE2, log_2_fold change=9.61, p = 0.016), and transcriptional activation (DNAJC2, log_2_fold change=12.6, p = 0.0051) in ZNF638 KD. C) ZNF638 KD in patient-derived GBM cell line results in upregulation of antiviral and immune pathways and programs. D) ZNF638 KD results in upregulation of several retrotransposons including LINE, LTR, and Alu elements (*P<0.05).

### Viral mimicry activates immune checkpoint blockade in GBM

To gain insight into the clinical impact of dsRNA expression on local tumor microenvironment, we employed multiplex immunofluorescence of naïve GBM, demonstrating an association between baseline dsRNA expression and both PD-L1 expression and CD8^+^ T-cell infiltration in patients treated with immunotherapy post-resection. **(Figure 7A, B, Supplementary Table 4)**

**Figure 7.**
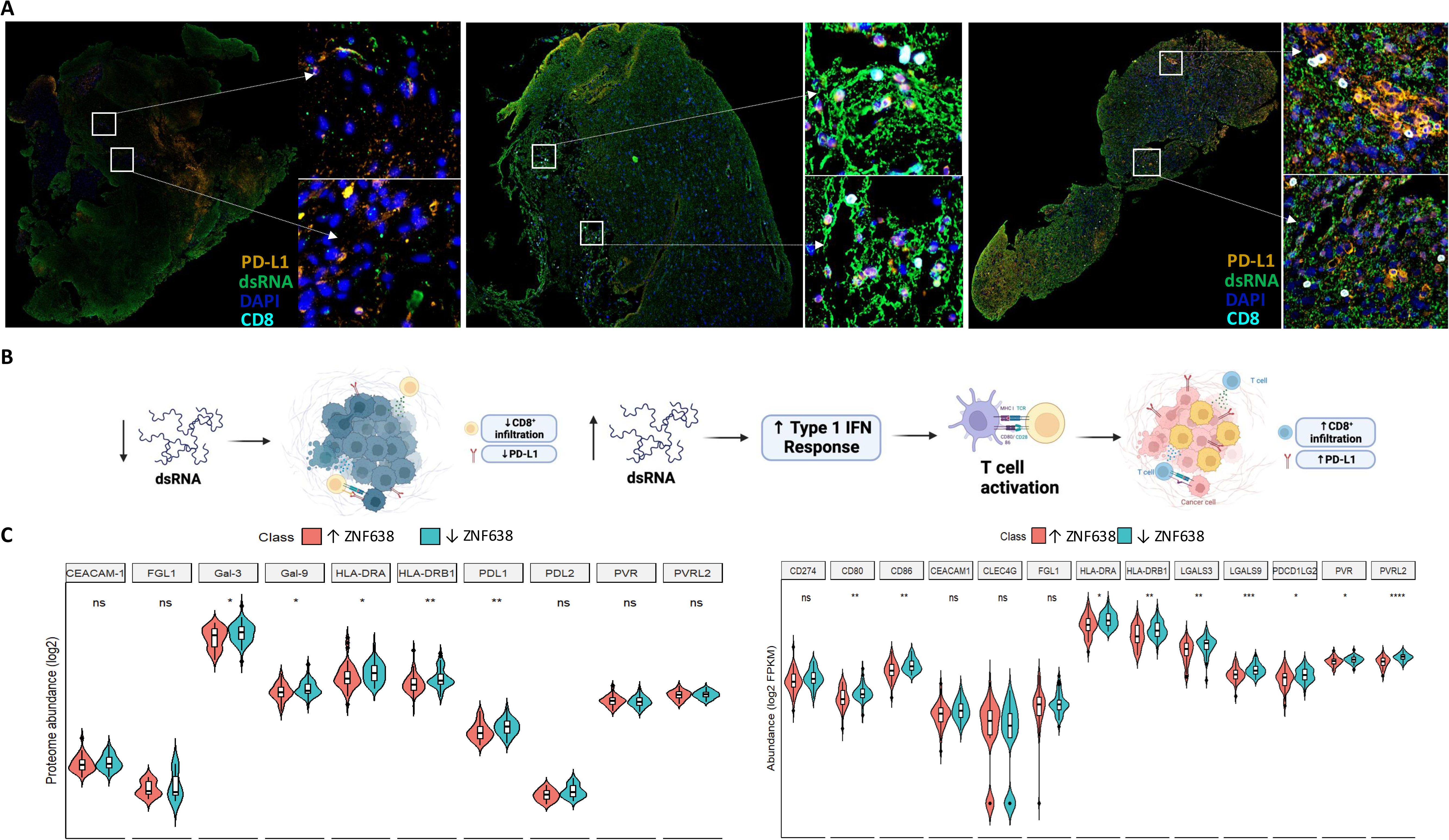
dsRNA expression is associated with increased checkpoint expression and CD8+ cell infiltration in high grade gliomas. A) Locoregional dsRNA expression is associated with increased PD-L1 levels and tumor infiltrating lymphocytes in human GBM. Yellow = PD-L1, Green = dsRNA, Blue = DAPI, Cyan = CD8. B) Increased expression of dsRNA stimulates a Type 1 IFN response to induce T-cell activation and infiltration and PD-L1 upregulation. Made in BioRender C) Proteomic and transcriptomic data obtained from the CPTAC data portal validate a negative relationship between expression of ZNF638 and immune checkpoint markers: PD-L1 (p<0.001), HLA-DRA (p<0.01), and HLA-DRB1 (p<0.001) (n_low_=50 and n_high_=50 in each group)

As previously demonstrated in lymphoma, melanoma, and colorectal cancer, activation of viral mimicry immune responses increases immune checkpoint presentation in solid tumors.^15,16^ Similarly, we discovered that ZNF638-KD significantly increased PD-L1 expression in multiple GBM cell lines (A172 and U87) via Western Blot and immunofluorescence. This suggests that dsRNA-sensing antiviral programs may potentiate immunotherapy in GBM. **(Figure 5A, C, Supplementary Figure 3A)** To corroborate the translational potential of our findings, we established stable shRNA knockdown of ZNF638 in murine GBM cell lines that have been previously validated to recapitulate the intrinsic immunosuppressive characteristics of GBM and its tumor microenvironment with poor basal checkpoint presentation.^29^ Importantly, ZNF638-KD in SB28 resulted in a substantial increase in expression of TLR3 and PD-L1 expression with attenuation of cellular proliferation. **(Supplementary Data Figure 5).** This relationship was corroborated in patient samples from the CPTAC data portal and TCGA RNA-seq datasets; high PD-L1 expression was significantly increased in patients with low ZNF638 expression. **(Fig. 7C)**

To further investigate the targetability of ZNF638 in sensitizing clinical responses to immune checkpoint inhibitors (ICI), we developed an immunocompetent syngeneic orthotopic model employing the SB28 cell line in C57BL/6 mice exposed to checkpoint immunotherapy **(Fig. 8A).** ZNF638-KD mice treated with intraperitoneal αPD-L1 survived significantly longer than CTL, ZNF638-KD alone, and sham vector + αPD-L1 groups. **(Fig. 8B)** This survival advantage was associated with tumor volumes 90-fold smaller compared to other experimental groups. **(Fig. 8C, Supplementary Figure 6A)**. Importantly, ZNF638-KD reduced intratumoral H3K9me3 and increased murine endogenous retroviral expression (RLTR6-M). **(Fig. 8D, Supplementary Figure 6A)**. Synergistic treatment with ZNF638KD and ICI significantly altered the GBM microenvironment by increasing inflammatory cytokine expression ( IL-2, IL-7, and IP-10, and interferons) and increasing expression of dsRNA sensing programs (RIG-I, TLR3). **(Fig. 8D,E, F).** Consistent with increased IP-10 levels, a chemotactic cytokine that attracts T-cells, ZNF638-KD tumor had enhanced CD8+ tumor-infiltrating lymphocytes (TILs) compared to other experimental groups. **(Supplementary Figure 6A).** Additionally, sera isolated from ZNF638-KD mice treated with ICI also revealed significant systemic elevation of IFN-alpha and IFN-γ levels with reduction of TNF-α levels. **(Figure 8F)** Using multiplex flow cytometry, we demonstrate that synergistic treatment increased populations of CD8+, CD4+ tumor infiltrating lymphocytes and a shift from monocytic myeloid derived suppressor cells (mMDSC) to granulocytic myeloid derived suppressor cells (gMDSC). While all MDSCs exhibit potent immunosuppressive activity in the tumor microenvironment, mMDSCs are known to have higher suppressive activity than gMDSCs.^24^ **(Supplementary Figure 6B).**

**Figure 8.**
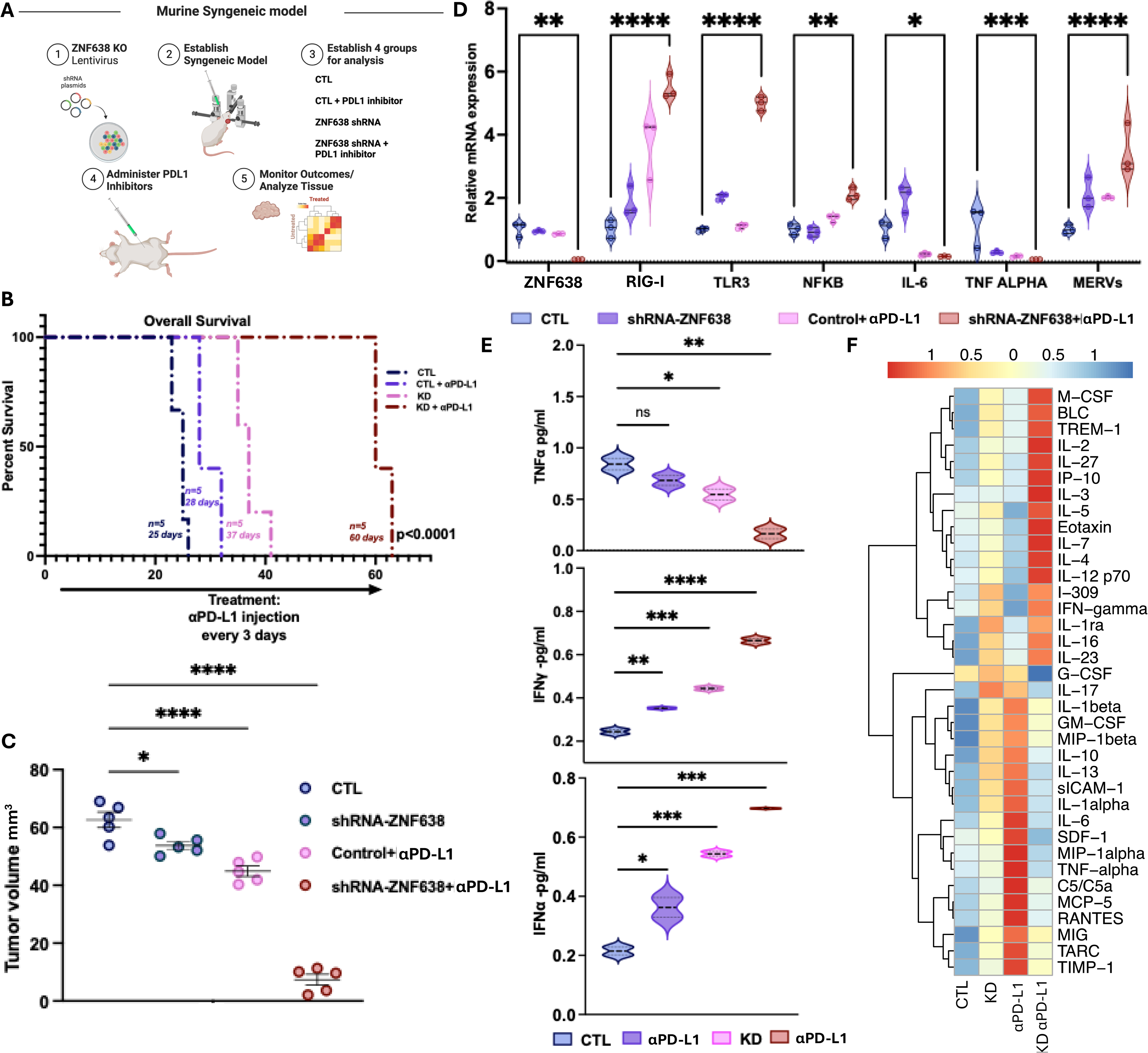
ZNF638 knockdown potentiates ICI response *in vivo*. A) Diagram of *in vivo* study design with syngeneic murine GBM model. Made in BioRender. B) Murine model with ZNF638 knockdown and PD-L1 inhibition demonstrates significantly improved survival relative to other treatment and control groups (n=5 per group, p<0.01). C) ZNF638 knockdown and PD-L1 inhibition significantly reduces tumor volume relative to all other groups (ANOVA, **** = p<0.00001, *** = p<0.0001, ** = p<0.001, * = p<0.01). D) ZNF638 knockdown + α-PD-L1 shows decreased expression of ZNF638 and increased expression of RIG-I, TLR3, and NFKβ transcripts as measured by qPCR. Additionally, there is significant upregulation of IL-6, TNF-α, and MERVs (RLTR6) (ANOVA, ****P < 0.00001, ***P < 0.0001, **P < 0.001). F) ZNF638 knockdown with PD-L1 inhibition significantly increased expression of IFNα and IFN-y as well as decreased expression of TNFα (one-way ANOVA, ****P < 0.0001, ***P < 0.001, **P < 0.01). G) Proteomic cytokine profiler array with hierarchical clustering depicts distinct cytokine profiles between all treatment/control groups with the greatest difference between CTL+ α PD-L1 mice and ZNF638 KD + α PD-L1 mice (one-way ANOVA *P<0.05).

### Viral Mimicry is associated with a clinical response to ICI in GBM

To gain insight into the clinical and molecular impact of dsRNA expression on the response to ICIs, we assessed temporal immunological changes in the GBM tumor microenvironment after a clinical response to ICI (nivolumab). In GBM patients, we noted that immune pseudoresponses were associated with elevated basal dsRNA expression and associated upregulation of PD-L1 expression and CD8+ cell infiltration. In ICI-naïve tumor specimens, the GBM tumor microenvironment remained relatively subdued with minimal infiltrating lymphocytes, low dsRNA expression, and low baseline PD-L1 expression. However, after initiation of ICI (anti-PD-1), new enhancement in the tumor cavity revealed histological pseudoprogression with substantial locoregional upregulation of PD-L1 expression in areas with elevated dsRNA expression. This immune pseudoprogression response was characterized by increased infiltration of CD8+ T-cells and activated microglia (IBA1) in regions enriched with high dsRNA expression. **(Figure 9A)** Given these findings, we sought to understand the role of ZNF638 in predicting long-term responses to immunotherapy in GBM using genomic and transcriptomic profiling of patients who received ICI (αPD-1, nivolumab or pembrolizumab).^30^ Consistent with our preclinical data, we discovered that ICI-Responsive GBM patients had significantly lower ZNF638 expression than non-responders, and was associated with a significantly improved overall survival. (**Figure 9B,C)** Importantly, this association is also upheld in other cancer types. In three separate cohorts of melanoma patients treated with Anti-PD-L1 or anti-CTLA4 therapy, low ZNF638 was significantly predictive of response. **(Figure 9B)** These results suggest that clinical responses to ICI are strongly correlated to innate antiviral immune signatures that could serve as clinical biomarkers for ICI.

**Figure 9.**
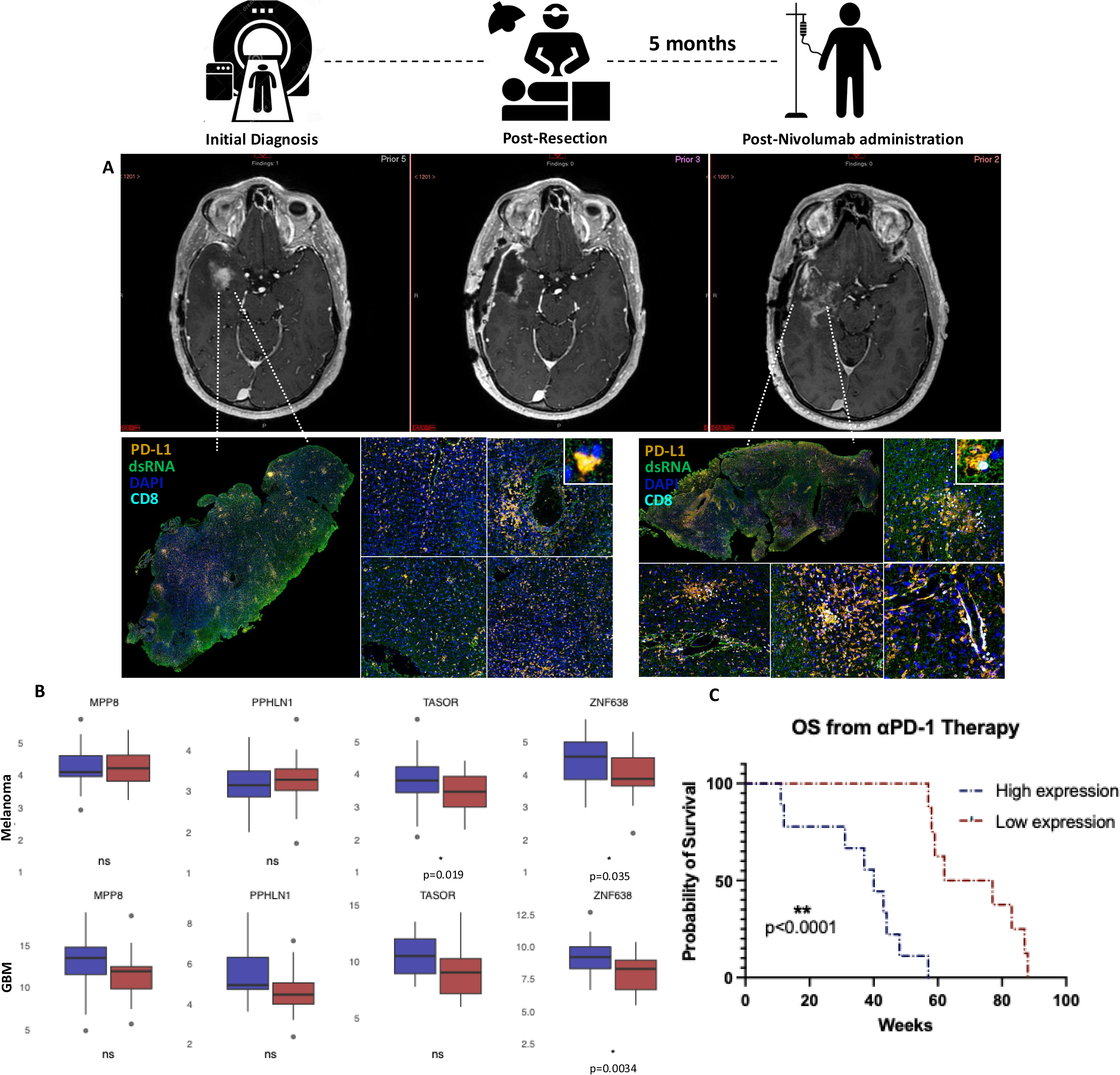
ZNF638 and dsRNA are biomarkers for ICI response in GBM. A) dsRNA expression correlates to increased CD8 T cell infiltration and PD-L1 expression in GBM patients receiving ICI. 60 y/o M with IDHwt rGBM with temporal contrast enhancing intra-axial tumor concerning for tumor recurrence (MR T1CE) with low baseline CD8 Infiltration (cyan) and PD-L1 expression (orange). Postoperative adjuvant ICI (anti-PD-1) resulted in increased enhancement in tumor cavity 5 months after surgical resection suggestive of immune pseudoprogression. Post-response multiplex immunofluorescence demonstrates increased dsRNA expression (green) associated with increased CD8 infiltration (cyan) and PD-L1 expression (orange) B): Clinical responders to ICI with rGBM (R=20, NR=18, median = 8.238 vs. 9.222 RPKM, p=0.0034) and melanoma (R=34, NR=49, median = 3.855 vs. 4.536 RPKM p=0.035) have markedly lower ZNF638 expression compared to non-responders to ICI C) Low ZNF638 expression portends improved survival in recurrent GBM receiving immunotherapy (PD-1 or PD-L1) (Mantel-Cox, *P<0.01).

## DISCUSSION

Glioblastoma is the the most common adult brain tumor with dismal prognosis despite current standard of care.^25^ Development of novel therapeutics, including immunotherapy for gliomas has largely failed due to its heterogenous and immunosuppressive tumor microenvironment.^26^ Therefore, treatment approaches that exploit immunosuppressive molecular marks are highly necessitated to improve therapeutic sensitivity for ICI. Gliomas are characterized by a heterogeneous profile of epigenetic dysregulation due to distinct methylation patterns (i.e. IDH1/2 mutations/CpG island methylator phenotype).^27,28^ Previously, we have demonstrated differential expression of retrotransposons in gliomas due to distinct locus-specific epigenetic regulation.^29–31^ Active transcription of these retroelements (REs) has been demonstrated to markedly improve sensitivity to checkpoint inhibition in colorectal cancer, lymphoma, melanoma, and renal cell carcinoma.^7–11,15,16^ Therefore, capitalizing on the expression of these REs in GBM via targeted epigenetic dysregulation may alter local and systemic immunogenicty and potentiate responses to ICI. Here, we have demonstrated that epigenetic activation of retrotransposon transcription through ZNF638 significantly alters tumor immunogenicity and improves survival via induction of viral mimicry.

The unique epigenetic state of these REs is controlled by the retroviral silencing complex, consisting of ZNF638, SETDB1, and the HUSH complex (MPP8, TASOR, PPHLN1). ZNF638 acts as the master regulator of this complex, responsible for recruiting the HUSH complex and mediating H3K9 trimethylation of retroviral DNA. Re-expression of these elements via widespread epigenetic dysregulation (DNMTis) has been demonstrated to induce dsRNA and dsDNA expression in multiple cancers.^15,32,33^ Other groups have shown that epigenetic reprogramming stimulates the innate anti-viral immune response via induction of the RIG-I signalling cascade.^34–36^ Our results established that targeted alteration of RE epigenetic control through ZNF638 knockdown induces RE-associated dsRNA expression by globally downregulating H3K9me3 and activating RIG-I/MDA5 pathway *in vitro* and in syngeneic GBM murine models.

Stimulation of the RIG-I/MDA5 pathway has been established as a critical element for responsiveness to immune checkpoint blockade (anti-CTLA-4 and anti-PD-1).^36^ This effect is mediated by Type 1 IFNs and upregulated IFNAR1.^34,36^ ZNF638 knockdown induced activity of RIG-I and its downstream effectors in syngeneic murine glioma models which resulted in increased PD-L1 expression. Importantly, our results demonstrated that ICI treatment significantly improved survival and reduced overall tumor growth in syngeneic GBM mouse models with ZNF638 knockdown. ZNF638 knockdown altered the GBM tumor microenvironment by enhancing immunogenicity through elevated Type 1 IFN responses and increased CD8^+^ T-cell infiltration. This antiviral immune response was conserved in patient-derived GBM neurospheres which exhibited upregulated immune and antiviral programs with concomitant global loss of genomic repressive marks. In line with these results, a recent study reported that GBM response to immunotherapy, and CD8 T-cell recognition, is associated with a MAPK-derived IFN-response phenotype by glioma cells.^37^

Previous studies have shown that expression of REs are directly associated with expression of Type I IFNs via direct binding of specific superfamilies (i.e. HERV-K) to intracellular dsRNA sensors.^34,38^ Russ et. al demonstrated that knockdown of MAVS, an associated downstream component of the dsRNA signaling cascade, subsequently reduces expression of Type 1 IFNs and related pro-inflamamtory cytokines.^38^ Our transcriptomic analysis of two independent glioma datasets further confirmed that ZNF638 expression is negatively correlated with expression of components of the dsRNA signalling pathway. Immune deconvolution of both independent datasets additionally demonstrated that ZNF638 expression is negatively correlated with estimated CD8^+^ and NK cell infiltration. Additionally, transcriptional activity of interferon-stimulated genes and pathways were shown to be inversely correlated with levels of ZNF638 expression. These findings point to the important role of the epigenetic regulators of REs on overall tumor immunogenicity and potential anti-tumor immune responses. Finally, a comprehensive evaluation of the landscape of retrotransposons demonstrated that multiple components of the retroviral silencing complex (PPHLN1 and TASOR) were widely negatively correlated with most HERV superfamilies. We also demonstrate with a thorough single-cell transcriptomic analysis that malignant cells lacking ZNF638 expressed higher levels of total retroelements, and were associated with diverse immune cellular profiles.

Overall, our results support a role for ZNF638 as a target to potentiate immune checkpoint inhibition through stimulation of the innate antiviral immune response. This finding was recapitulated longitudinally in the local tumor microenvironment (TME) of GBM responders to ICI. Temporal alterations in the local GBM TME in response to αPD-1 immunotherapy were associated with diffuse dsRNA expression and increases in both PD-L1 expression and CD8^+^ T-cell infiltration. More importantly, ZNF638 expression was a biomarker of clinical response to immunotherapy across multiple tumor types including rGBM and melanoma. Therefore, eliciting dsRNA expression may be a therapeutic modality to potentiate the efficacy of ICI in GBM

Clinical translation of viral mimicry for GBM has been proposed previously through epigenetic reprogramming. Viral mimicry and cytosolic dsRNA expression can be induced through radiation, epigenetic drugs such as DNMTIs or HDACs, and synthetic molecules such as RIG-I agonists. ^39,40^ However, the systemic toxicity of epigenetic therapies and local toxicity of radiation have limited their clinical translation in oncology.^41,42^ Synergistic treatments using epigenetic reprogramming and ICI are currently underway in a variety of solid tumors, and have shown some promise in early phase clinical trials. ^43–46^

Although viral mimicry has been shown to enhance immunotherapy responses in cancer, it is critical to consider the pleiotropic role of REs in the GBM TME. We have previously demonstrated that expression of certain HERV-K families (HML-2, HML-6) are associated with synthesis of full-length retroviral proteins that may contribute to an oncogenic phenotype and tumor stemness.^29,30^ In our analysis, targeting ZNF638 did not affect endogenous HERV-K expression, suggesting differential control of older viral dsRNA elements and more recently integrated HERV-K loci. Therefore, we suspect that there remains a balance between anti-viral immune programs (overall RE expression) and specific oncogenic viral programs (HERV-K). Further RE-specific methylation profiling may elucidate the specific retroelements associated with viral mimicry immune responses.

Overall, ZNF638 is a biomarker of clinical response to ICI in GBM, suggesting that ZNF638 expression could not only predict clinical responses but may also serve as a target to potentiate immunotherapy across multiple tumor types. It is unknown how ZNF638 may interact with other markers of response to immunotherapy, such as clonal tumor mutational burden, STING activation, DNA replication stress, and MAPK signaling. Future studies are needed to clarify these potential interactions.

Taken together, there may be a clear role for our findings as an adjuvant therapy to both enhance anti-tumor innate immune responses and potentiate immunotherapy. Overall, these results inform the direct impact of ZNF638 on mediating viral mimicry immune responses in GBM, uncovering an avenue for clinical translation of epigenetic therapies that leverage the retroviral landscape of GBM.

## METHODS

### Cell culture and Transfection

Neurospheres derived from patient GBM samples (GBM43, IDH WT, 69-year-old male) were sourced from the Mayo Clinic Brain Tumor Patient-derived Xenograft National Resource.^47^ These were cultured in DMEM/F12 with GlutaMAX (Invitrogen), supplemented with 10 ng/mL epidermal growth factor, fibroblast growth factor, B27, N2 (Invitrogen), 1% penicillin-streptomycin, Heparin, and 1% sodium pyruvate (Fisher Scientific). Established cell lines (A172, U87, and normal human astrocytes) were acquired from ATCC and maintained per the manufacturer’s guidelines in their respective media: A172/U87 in DMEM with 5% FBS and penicillin/streptomycin, the murine SB28 cell line (gifted by Dr. Defne Bayik, University of Miami) in RPMI with 5% FBS and penicillin/streptomycin, and normal human astrocytes in Astrocyte Growth Medium Bullet Kit (Lonza). Neurospheres and established cell lines were dissociated using TrypLE Express (Invitrogen) and cells were not used beyond passage number 20. For ZNF638 knockdown experiments (Thermo Scientific, AM16708), predesigned siRNAs and scrambled negative control siRNA were utilized. Transfections of siRNA duplexes into A172, U87, and GBM43 cell lines were performed using Lipofectamine RNAiMAX (Thermo Scientific, 13778150) following the manufacturer’s instructions. Post-transfection, RNA and protein were extracted for downstream analyses including RNA sequencing, ATAC sequencing, western blotting, and immunohistochemistry.

### Western blot analysis

Cells were treated with ZNF638 siRNA and cultured for 72 hours prior to protein collection. Total protein was extracted using RIPA buffer (Sigma-Aldrich) with protease/phosphatase inhibitors (ABCAM, Waltham, MA). Protein concentrations were determined using Bio-Rad Protein Reagents, following the manufacturer’s protocol. The protein samples were denatured in a mixture of SDSPAGE Reducing Agent (×10) and RIPA buffer (×1), separated on a 4%–20% Tris-Glycine SDS-PAGE gel (Bio-Rad), and transferred to nitrocellulose membranes using the iBlot transfer device (Thermo Fisher Scientific, Waltham, MA). Membranes were blocked with 5% Blotting-Grade blocker in TBS (Bio-Rad) with Tween (Thermo Fisher Scientific), followed by incubation with primary antibodies overnight. After washing with TBS-T, membranes were probed with either anti-rabbit or anti-mouse secondary antibodies (Invitrogen) and visualized using ECL chemiluminescence (Thermo Fisher Scientific). Detailed information about the specific antibodies used is provided in Supplementary Table 5. For histone analysis, histones from A172 cells treated with scramble and ZNF638 siRNA were purified using a histone extraction kit (Active Motif) as per the manufacturer’s instructions. Twenty micrograms of each lysate were separated on a gel and probed for total H3, H3K9me3, and H3K9me2.

### Spatial Transcriptomics

Raw counts from the spatial transcriptomics experiment were exported from Nanostring Geomx into R studio version 4.4.1. The Bioconducter package *GeomxTools* was used for quality control and data filtering. The R package *dplyr* was also used for data manipulation. Regions of interest were sorted into high and low levels of ZNF638 expression. Two regions from one glioblastoma patient were analyzed with heterogeneous ZNF638 expression. Expression data was sorted by pathway, using gene sets from the *MSigDB* R package. This was visualized with a radial plot using R package *ggplot2*.

### Analysis of ZNF638 and REs in Transcriptional States and Cell Types

Single-cell RNA sequencing data obtained from Johnson et. al^22^ was annotated using a custom bioinformatics pipeline for retrotransposon counts from TE-transcripts.^20^(M. Hammel Laboratory, Cold Spring Harbor Laboratory, Cold Spring Harbor, New York, USA) This data was analyzed using standardized workflow for object creation, integration, normalization, and visualization in Seurat in RStudio.^48,49^ Cells were filtered based on percentage of mitochondrial transcript expression and number of detected features. Filtered data were normalized and scaled using inherent Seurat functions. Cell types and transcriptional states were annotated based on established cell-type and cell-state specific markers.^23^ A custom script was developed for scoring transcriptional state signatures based on individual cells. Ranked lists were used as input to score each cell for cell type and transcriptional state signatures. These were visualized using DimPlot. The inherent Seurat function FindMarkers was used to calculate log_2_fold change and adjusted P-values for unique markers of each cluster. These were used to verify cell type and transcriptional states of clusters labelled by ranked list input. All UMAP plots were generated using Seurat functions RunUMap and DimPlot. All heatmaps and feature plots were visualized using Seurat functions DoHeatmap and FeaturePlot respectively.

### Preparation of cDNA from Tumor Tissue and Cell Lines

RNA was isolated from tumor tissue and cell line samples using the Trizol extraction protocol. Quantification of RNA was performed using the NanoDrop 2000 (Thermo Fisher Scientific), and the concentration was adjusted to 1000 ng/μL. The RNA was then reverse transcribed using the iScript™ cDNA Synthesis Kit (1708890). Quantitative PCR (qPCR) was employed to amplify and detect target transcripts on the cDNAs, utilizing 5 μM primers. Detailed information about the specific antibodies used is provided in Supplementary Table 6.

The qPCR cycling conditions were set as follows: initial denaturation at 95°C for 20 seconds, followed by 35 cycles of 95°C for 20 seconds and 60°C for 30 seconds, using the Fast SYBR Green Master Mix. Normalized CT values were used to quantify the expression of target transcripts through the ΔΔCT method. All qPCR runs included reverse transcription negative controls, which did not show amplification. Each qPCR experiment was conducted in both biological and technical triplicates.

### Immunofluorescence

Cells were cultured as previously described and treated with ZNF638 siRNA for 6 hours. Forty-eight hours post-treatment, cells were seeded into 2-chambered slides at a density of 2.5 x 10^6^ cells/cm^2^. The cells were then fixed with 4% paraformaldehyde (Thermo Fisher Scientific) for 20 minutes and washed with PBS. For membrane permeabilization, cells were incubated with 0.01% Triton X for 5 minutes, followed by PBS washes and blocking with 10% normal goat serum. Primary antibodies (diluted 1:200) were applied overnight, then washed with PBS. Subsequently, slides were incubated with fluorescent-tagged goat anti-rabbit and goat anti-mouse secondary antibodies in 2% normal goat serum. Detailed information about the specific antibodies used is provided in Supplementary Table 5. For control, control + αPDL-1, shRNA-ZNF638, and shRNA-ZNF638 + αPDL-1 samples, slides were deparaffinized and rehydrated using xylene and graded ethanol (100%, 95%, 70%). Antigen retrieval was performed by steaming slides in citrate buffer for 20 minutes. Permeabilization, blocking, and primary/secondary antibody incubations were conducted as described above. Images were captured using the EVOS M700 microscope.

### RNA sequencing

Ribosomal depleted RNA was extracted from patient derived neurosphere cell lysate and sequenced using a lncRNA prep (250-300 bp, paired-end, 50 million reads). RNA-sequencing was performed by Novogene (Sacramento, CA) using the Illumina NovaSeq 6000 PE150 platform. FastQ files were prepared and aligned to the human Hg38 reference genome, and subjected to external quality control. A custom bioinformatics pipeline from TE-Transcripts.(M. Hammel Laboratory, Cold Spring Harbor Laboratory, Cold Spring Harbor, New York, USA) was used to annotate retrotransposon counts^20^ Differential expression analysis was conducted using DESeq2 with multiple testing corrections. Functional pathway and gene ontology analysis for each biological condition was conducted using “clusterProfiler” from Bioconductor. All plots were visualized in R using “ggplot” or Python using “seaborn”.

### ATAC-Sequencing

The preparation and sequencing of libraries for ATAC-seq were performed by Novogene (Sacramento, CA). Paired-end sequencing was performed using the Illumina NovaSeq 6000 PE150 platform. FastQ files were aligned to the human Hg38 reference genome. Differentially accessible peaks were identified after DESeq2 normalization. Integrative Genomics Viewer was used to visualize tracks (version 2.17.4, Broad Institute).^50^ Functional pathway and gene ontology analysis for each biological condition was conducted using “clusterProfiler” from Bioconductor.

### Proliferation assay

shRNA-NC or shRNA-ZNF638 GBM cells were seeded at 1,000 cells/well in a 96-well E-plate in biological and technical triplicates. Cell proliferation was measured with xCELLigence RTCA DP instrument according to the manufacturer’s instructions (ACEA Bioscience, CA, USA) and visualized for over 7 days in culture or until cell proliferation plateaued.

### Intracranial orthotopic xenografts

The mouse GBM cell line (SB-28) cell line was transduced with either lentivirus CTL shRNA and ZNF638 shRNA with an mCherry tag were cultured for 72 hours and sorted using BD FACS Aria (BD Biosciences). The sorted mcherry-positive cells were maintained in cell culture for 3 days. The harvested cells were used for implantation. 2 × 10^4^ cells resuspended in 3 μL PBS and implanted into the right frontal lobe of Immunocompetent six-week-old C57BL/6 female mice were obtained from The Jackson Laboratory using the following coordinates (AP, 1.5 mm; DV, 3 mm; ML, 2 mm). After tumor establishment on day 7, mice were treated with an anti PD-L1 (Bio cell) via intra-peritoneal injection for every 3 days until the survival of the mice. Tumor volumes were measured and calculated twice per week using the modified ellipsoid formula ½ × (length × width2). Brain tumors were harvested immediately after sacrifice, fixed with 4% paraformaldehyde, and sectioned for histopathology. Whole blood was collected and allowed to clot at room temperature, then the serum was isolated by centrifuging at 1,000-2,000 x g for 10 minutes. The resulting supernatant and tumor tissues were preserved for downstream analysis, including ELISA and qPCR. All animal experiments were approved by the Institutional Animal Care and Use Committee at the University of Miami and performed in accordance with the guidelines.

### Immune profiling

For immune profiling, the tumor tissue was digested using previously described methods from Newton et al., 2018.^51^ In brief tumors were removed from adjacent brain using microdissection. Tumour tissue were mechanically dissociated on a 40 μm strainer and washed with PBS before transferring into 96-well round-bottom plates (Thermo Fisher Scientific). Samples were stained with 1:1,000 diluted LIVE/DEAD Fixable Stains (BioLegend) in PBS for 10 minutes on ice. Following a wash step, cells were resuspended in FcR Blocking Reagent (Miltenyi Biotec) at a 1:25 dilution in PBS/2% BSA (Sigma-Aldrich) for 10 minutes on ice. Fluorophore-conjugated antibodies diluted 1:50 were added to suspensions, and cells were further incubated for 20 minutes on ice. A list of antibodies can be found in Supplemental Table 5. Samples were acquired with Cytek Aurora (Cytek Biosciences) and analyzed using FlowJo (v10.7.2, BD Biosciences).

### Cytokine Array Panel

Tumor samples were excised from sacrificed mice and homogenized in PBS supplemented with protease inhibitor and Triton-X. Proteins were isolated and cytokines were measured using Mouse Cytokine Array Panel (ARY006, R&D Biosystems. Membranes were imaged using ECL chemiluminescence (R&D Biosystems) and data analysis was performed using HLImage++ 6.2 software.

### Immunohistochemistry

The immunohistochemistry (IHC) detection kit was purchased from Abcam (ab64264). Experiments were performed according to the manufacturer’s instructions. Anti-H3K9Me3(Cell Signaling D4W1U) and anti-CD8 (Cell Signaling D4W2Z) were diluted 1:200. Images were captured using EVOS M700 microscope at 10X and 20X magnification.

### Tissue Microarray

Unstained tissue microarray slides (core size 1.0 mm) were procured from Tissue Microarray (GL2082a), encompassing brain tumor and brain tissue samples, including 37 GBM and 14 normal cerebrum, with duplicate cores per case. The slides were processed using an IHC detection kit from Abcam (ab64264), following the manufacturer’s instructions, with the ZNF688 antibody at a 1:200 dilution. Immunostained slides were scanned using a High-throughput VS120 Olympus slide scanner with x4-x40 air lenses. The scanned images were then analyzed in a blinded fashion using Qupath image viewer software (v0.5.0-Mac-x64), with curated regions of interest outlined manually.

### RNA Immunoprecipitation qPCR

A172 cells treated with either scramble or ZNF628 siRNA (5.0 × 10^6 cells) were harvested, and cytoplasmic fractions were extracted using the Nuclear Extract Kit (Active Motif; no. 40010) according to the manufacturer’s instructions. To isolate RNA, an equal volume of 70% ethanol was added to the cytoplasmic fractions, and RNA purification was performed using the RNeasy Plus Mini Kit (Qiagen; no. 74106) following the manufacturer’s protocol. The total RNA was dissolved in 38 μL RNase-free water. Two microliters of total RNA were used as input, and the remaining RNA was divided into two tubes.

For each RNA immunoprecipitation pulldown, 2 μg of J2 antibody (SCICONS; no. 10010200) and mouse control IgG2a (Abcam; no. ab18413) were conjugated to 20 μL protein G agarose (Millipore; no. 16-266) by rotating overnight at 4°C. To digest single-stranded RNA, 1 μL of RNase A (Sigma-Aldrich; no. R6513) was added to each tube, and then mixed with 1 mL IP buffer (50 mmol/L Tris-HCl [pH 7.4], 125 mmol/L NaCl, 1 mmol/L EDTA, 0.1% Triton X-100). The RNA samples were incubated with antibody-conjugated protein G agarose beads overnight at 4°C. Beads were washed three times with IP buffer and then incubated in 50 μL proteinase K digestion solution (1× TE, 100 mmol/L NaCl, 1% SDS, and 1 μL of 20 mg/mL Proteinase K solution (Thermo Fisher Scientific; no. AM2546)) for 20 minutes at 45°C to isolate RNA. After centrifugation, 50 μL of the supernatant was added to 300 μL Buffer RLT Plus from the RNeasy Plus Mini Kit (Qiagen; no. 74106) to purify the RNA. The final product containing dsRNA was denatured for 5 minutes at 95°C, followed by reverse transcription using qScript cDNA SuperMix (iScript™ cDNA Synthesis Kit, 1708890). qRT-PCR was then performed using the primers listed in Supplementary Table 5 with the CFX96 Touch Real-Time PCR Detection System.

### Flow Cytometry

GBM43 neurospheres treated with scramble or ZNF638 siRNA were trypsinized and fixed using the CytoPerm/Fix kit (BD). The cells were then stained overnight at 4°C with mouse anti-dsRNA antibody (Scicons). Following staining, the cells were washed and resuspended in FACS buffer (1% FBS, 0.9% sodium azide in PBS). Flow cytometry analysis was performed using a BD LSRFortessa Cell Analyzer. Up to 50,000 cells were collected and gated to exclude non-viable cells.

### Co-Immunoprecipitation

For immunoprecipitation, the A172 scramble and ZNF638 siRNA cells were lysed by RIPA lysis buffer using Immunoprecipitation kit (ab206996) containing protease inhibitors. Lysates were extracted and incubated with protein A/G agarose for preclearing, then immunoprecipitated with the primary antibody (ZNF638) overnight followed by incubation with protein A/G agarose for 2.5 h. The precipitates were eluted and subjected to SDS gels for Co-IP assay using SETDB1, TASOR and MMP8 antibodies.

### Enzyme-Linked Immunosorbent Assay (ELISA)

Serum samples were collected and proceeded for ELISA following manufacturer protocol using IFN-γ (MIF00-1, R&D Biosystems), TNF-alpha (MTA00B-, R&D Biosystems), and IFN-alpha (42120-1, R&D Biosystems). Absorbance in each well was then read at 450 nm using an Spectramax M5 automated reader (Molecular Devices, San Jose, CA), and evaluated using system-associated Softmax pro software v4.8.

### Multiplex Immunofluorescence

Paraffin-embedded tissue slides underwent deparaffinization and permeabilization through washes with xylene and ethanol. Antigen retrieval was performed using a 10 mM sodium citrate buffer (pH 6.0), heated for 2 minutes in an 800 W microwave (GE model PEM31DFWW) at full power. Following antigen unmasking, the slides were blocked using Background Buster (Innovex Biosciences, NB306) and FcR blocking solution (Innovex Biosciences, NB309). Next, tissue sections were incubated at room temperature for 60 minutes with an 11-plex primary antibody cocktail, followed by multiple washes (three times with dH2O). This was followed by secondary antibody incubation, with sections washed again (three times with PBS and three times with dH2O). Subsequently, the tissue sections were counterstained with 1 μg/mL DAPI (Thermo Fisher Scientific) for pixel-pixel registration reference. Imaging was conducted using an Axio Imager.Z2 scanning fluorescence microscope (Carl Zeiss) equipped with a 20×, 0.8 NA Plan-Apochromat (Phase-2) nonimmersion objective (Carl Zeiss) and a 16-bit ORCA-Flash 4.0 sCMOS digital camera (Hamamatsu Photonics). Each labeling antibody was captured sequentially at its specific wavelength and digitized individually using ZEN2 imaging software (16-bit). Channels for antibodies of interest (dsRNA[green], PD-L1[yellow], CD8+[cyan], and DAPI[blue]) were compared between samples qualitatively. Patient characteristics and pathologies are described in **Supplementary Table 4.**

### Human GBM ICI Response and Survival Analysis

RNA count matrices and survival data were obtained from Zhao et. al 2019^52^ on a cohort of 38 patients treated with Anti-PD1 immunotherapy. Survival and response data on 3 cohorts of Melanoma patients treated with Anti-PD1 therapy or CTLA-4 blockade obtained from Van Allen et. al 2015, Snyder et. al 2015, and Hugo et. al. 2017.^53–55^ Integration methods are described by Noviello et. al 2023.^56^ Boxplots for responders and nonresponders in both GBM and melanoma cohorts were generated in R using ggplot. Kaplan-Meier survival analysis was calculated and visualized in PRISM. Statistics are detailed below and in **Supplementary Table 7.**

*Statistics.* All experiments were conducted with biological replicates or triplicates and confirmed with technical triplicates. Data are shown as mean ± SEM. Full details for statistics are detailed within each subsection and within Supplementary Table 7. Tests used include unpaired 2-tailed *t* test, Mann-Whitney test, 1-way ANOVA, and log-rank (Kaplan-Meier). *P* values of less than 0.05 were considered significant.

*Data Availability.* All de-identified data will be made availalble from the corresponding author upon request and signature of data transfer agreement.

*Study Approval*. All animal experiments were approved by the Institutional Animal Care and Use Committee at the University of Miami and performed in accordance with the guidelines (protocol #22-117-adm01).

## AUTHOR CONTRIBUTIONS

JC, DS, and AHS were involved in study initiation and design. JC, DS, JD, CKR, JCR, AJH, VG, YZ, AMS, MMV, DM, VG, SRR, VL, AA, RT, NS, CD, KJ, MDLF, RA, TN, MEI, RJK, AI, AN, JH, MC, KC, MF, DB, AHS contributed to data collection, analysis, and running experiments. JC prepared the manuscript, which was reviewed and approved by all authors. AHS supervised all aspects of this work.

